# Exposure to oxacillin subinhibitory concentrations and *in vitro* induction of resistance expression in heteroresistant and non-heteroresistant oxacillin-susceptible *mecA*-positive *Staphylococcus aureus* (OS-MRSA)

**DOI:** 10.1101/2023.01.24.521449

**Authors:** Denise Braga Schimidt, Helvécio Cardoso Corrêa Póvoa

**Affiliations:** Department of Microbiology and Parasitology, Biomedical Institute, Federal Fluminense University (UFF), Niterói, Rio de Janeiro, Brazil; Department of Microbiology, Immunology and Parasitology, Faculty of Medical Sciences, Rio de Janeiro State University (UERJ), Rio de Janeiro, Brazil; Department of Basic Sciences, Federal Fluminense University (UFF), Nova Friburgo Health Institute, Nova Friburgo, Rio de Janeiro, Brazil

## Abstract

Resistance expression can occur as a consequence of irrational use of antibiotics, since this implies dissemination of subinhibitory concentration (sub-MICs) in different environments and generates a selective pressure. The study aimed to evaluate the *in vitro* influence of selective pressure on antibiotic resistance expression in two oxacillin-susceptible *mecA*-positive *Staphylococcus aureus* (OS-MRSA), through exposure to oxacillin sub-MICs. One heteroresistant OS-MRSA strain and a non-heteroresistant strain, both isolated from nasal colonization were exposed to two-fold serial dilutions of oxacillin (0,125 a 256 μg/mL) during seven consecutive days. Disc diffusion test was used to determine the susceptibility to several antibiotics and population analysis profile (PAP) was used to evaluate the expression of oxacillin resistance before and after antibiotic exposure. Susceptibility to non-β-lactam antibiotics was not altered but changes in phenotypic expression of penicillin and oxacillin resistance were observed. Both OS-MRSA strains began to express homoresistance (oxacillin MIC = 256 μg/mL) and had no penicillin zone inhibition after induction, different from that was observed before oxacillin exposure, which suggested increased in β-lactamase production. *In vitro* selective pressure with oxacillin stimulated β-lactamase production and led phenotypic expression of oxacillin resistance in heteroresistant and non-heteroresistant OS-MRSA, which became homoresistant. This reinforces the impact that irrational use of antibiotics has on individuals colonized by *S. aureus* and on the population, emphasizing that the emergence and spread of resistance to antibiotics represent a process of evolution in response to selective antimicrobial pressure.

## Introduction

Antibiotics are among the most commonly prescribed drugs in clinical practice. However, most prescriptions (30 to 50%) are unnecessary due to errors in treatment indication, incorrect choice of antibiotics and duration of therapy [1]. Misuse and overuse are frequently attributed to the development of resistance because leads to the spread of different antibiotic concentrations, including subinhibitory concentrations (sub-MICs), in human, animals and in the environment. This results in antimicrobial selective pressure that acts in commensal and pathogenic bacteria [2], favoring the genetic evolution of microorganisms and the emergence and spread of antibiotic resistance [3]

In *Staphylococcus* aureus, oxacillin resistance is mainly related to expression of a modified PBP with low β-lactams affinity (PBP2a), which is encoded by the *mecA* gene. However, there are strains known as OS-MRSA (oxacillin-susceptible *mecA*-positive *Staphylococcus aureus*), that have *mecA* but are phenotypically susceptible to oxacillin [4, 5].

Mechanisms involving the existence of strains with this profile had not yet been clarified. But *mecA* presence indicates that this gene may be transcribed, leading to phenotypic resistance expression, if there are conditions able to inducing its transcription. One example is exposure to β-lactams antibiotics [6].

Sub-MICs of β-lactams have been shown to be able to inducing *in vitro* the expression of oxacillin resistance in OS-MRSA strains [5, 7–12]. Therefore, the use of these antibiotics to treat OS-MRSA infectious may result in change of susceptibility to oxacillin, resulting in therapeutic failure [4, 13]. However, little is known about the proportion of strains that convert to resistance phenotype [12].

OS-MRSA strains have already been found colonizing the anterior nares [8, 14] and these might be constantly stimulated to express β-lactam resistance because are frequently exposed to antibiotics sub-MICs, due to irrational and empirical use of antibiotics, as well as inadequate prescription. This study aimed to investigate effects that the *in vitro* exposure of commensal nasal OS-MRSA strains to sub-MICs of oxacillin has on one of the most important factors for the success of *S. aureus* as an opportunistic pathogen: the resistance expression to antibiotics.

## Material and methods

### Bacterial strains

Two OS-MRSA penicillin-resistant strain (SA607 and SA786), isolated in February 2014 from nasal colonization swab of hospitalized pacients was studied. A methicillin-susceptible *S. aureus* (MSSA) strain (SA292) also isolated from anterior nares was used for comparison. This strain was penicillin-susceptible but β-lactamase positive, according to penicillin disk diffusion zone-edge test [15]. In addition, *S. aureus* ATCC 25923 (MSSA, *mecA* negative, β-lactamase negative) and *S. aureus* ATCC 33591 (MRSA, *mecA* positive) were used as controls.

*Staphylococcus aureus* was isolated in pure culture using Müller-Hinton (MH) agar (Difco^®^, MD, USA) and were identified through Gram stain, catalase, tube coagulase (Coagu-Plasma, Laborclin^®^, PR, Brazil) and DNase (Himedia^®^, Mumbai, India) tests. Phenotypic identification was confirmed by detection of a *S. aureus* specific DNA fragment by PCR [16] and through MALDI-TOF MS using Autoflex Speed (Bruker Daltonic^®^, Billerica, EUA) and the MALDI Biotyper software package version 3.1. Both strains had scores > 2.0 which means reliable identification at species level.

### Antibiotic susceptibility

Antibiotic susceptibility was determined by disc diffusion method and broth microdilution according to CLSI guidelines [15]. Bacterial suspensions were prepared from isolated colonies, obtaining an OD of 0.17 at 520 nm, which correspond to approximately 10^8^ CFU/mL (0.5 McFarland standard).

The disc diffusion method was performed with penicillin (10U), oxacillin (1μg), cefoxitin (30 μg), gentamycin (10 μg), erythromycin (15 μg), tetracycline (30 μg), ciprofloxacin (5 μg), nitrofurantoin (300 μg), clindamycin (2 μg), sulfamethoxazole-trimethoprim (23,75 μg/1,25 μg), chloramphenicol (30 μg) and rifampicin (5 μg) (Sensifar, Cefar^®^, SP, Brazil).

Broth microdilution was performed to determine oxacillin MIC, with 5 x 10^5^ CFU/mL. However, the oxacillin susceptibility was determined by the disc diffusion test with cefoxitin, as recommended by CLSI [15].

### Population Analysis

The evaluation of the phenotypic expression of oxacillin resistance was performed through population analysis profile (PAP), according to Ikonomidis *et al* [17] Overnight growth was used to prepare bacterial suspensions that were adjusted to a density equivalent to a 0.5 McFarland standard (~ 10^8^ CFU/mL). From this starting suspension, 100 μL (approximately 10^7^ CFU) were spread onto MH agar plates containing oxacillin (0.125 to 256 μg/mL). After 48 hours of incubation at 35 °C the number of colony forming units (CFU) was counted.

The interpretation followed the recommendations of El-Halfawy & Valvano [18] considering the difference between the lowest concentration with maximum inhibition of growth and the highest non-inhibitory concentration. When the difference was ≤ 4-fold, the population was classified as susceptible and homogenous; equal to 8-fold, was intermediate heteroresistant; and > 8-fold, was heteroresistant.

### Molecular characterization

The release of DNA from overnight cultures was done by a thermal lysis method [19]. The DNA obtained was used as template for detection of *mecA* and *lukS-PV/lukF-PV* genes and for typing *SCCmec* by PCR. The presence of *mecA* and PVL genes was performed as previously described by Zhang *et al* [20] and von Eiff, Friedrich, Becker [21], respectively. *SCCmec* typing was determined by a multiplex PCR [20] with primers for *SCCmec* types and subtypes I, II, III, IVa, IVb, IVc e IVd.

### Oxacillin induction

Serial passage experiments were carried out as described by Kampf *et al* [8], with some modifications. Daily exposure to oxacillin was performed by broth microdilution using two-fold oxacillin serial dilutions (0.125 to 256 μg/mL) in Müller-Hinton broth (Difco). On the first day of exposure 5 × 10^5^ CFU/mL of each strain were inoculated in 100 μL of oxacillin dilutions. Plates were incubated aerobically at 35 °C during 48 hours to allow initial adaptation of bacteria to the environmental stress [9].

The next passage was inoculated on the same oxacillin concentrations (0.125 to 256 mg/L) with a 1:100 dilution of the vial with a highest concentration of oxacillin and visible growth from previous passage. Plates were incubated aerobically at 35 °C during 24 hours. This process was repeated for up to ten days. Three independent experiments were conducted in triplicates.

After induction strains were again submitted to population analysis (PAP) with oxacillin and to disc diffusion with the same antibiotics used before oxacillin exposure.

## Results

### Antibiotic susceptibility and genotypic characterization before induction

SA607 and SA786 had oxacillin MICs equal to 0.5 μg/mL and 0.25 μg/mL, respectively. Both were susceptible to all antibiotics tested, with the exception of penicillin. Therefore, strains were classified as oxacillin-susceptible according to oxacillin MIC (< 2 μg/mL) and cefoxitin disc diffusion (zone diameter > 22 mm) using the criteria of CLSI [15]. PCR was positive for the *mecA* gene. SA607 carried *SCCmec* type IVa as well as *lukS-PV/lukF-PV* (Panton-Valentine leukocidin – PVL genes) while SA786 did not have the PVL genes and the *SCCmec* was not typable (Tab. 1).

**Table 1.** Identification, antibiotic susceptibility and molecular characterization of strains.

|  | Species Identification |  | Antibiotic susceptibility<sup>a</sup> |  | Molecular characterization |  |  |
| --- | --- | --- | --- | --- | --- | --- | --- |
|  | MALDI-TOF (score) | PCR (S. aureus) | Disc diffusion (resistance) | Oxacillin MIC (µg/mL) | *mecA* | *SCCmec* | PVL |
| SA29<br>2 | 2.245 | + | ERY,<br>CLI<sup>b</sup> | < 0.125 | - | - | - |
| SA60<br>7 | 2.318 | + | PEN | 0.5 | + | IVa | + |
| SA78<br>6 | 2.278 | + | PEN | 0,25 | + | - | - |

<sup>a</sup>all strains were susceptible to the others antibiotics tested. <sup>b</sup>inducible clindamycin resistance. ERY: erythromycin; CLI: clindamycin; PEN: penicillin.

### Oxacillin induction and antibiotic susceptibility after induction

Exposure to sub-MICs of oxacillin for five to six days led OS-MRSA to express high and homogeneous oxacillin resistance (MIC of 256 μg/mL). Before six days both had already altered the expression of oxacillin resistance. From the first to the second day, SA607 changed the susceptibility profile to the resistance and MIC increased 128-fold. SA786 only became resistant at the fourth day of exposure, with an increase of 256-fold in MIC (Fig. 1).

**Figure 1.**
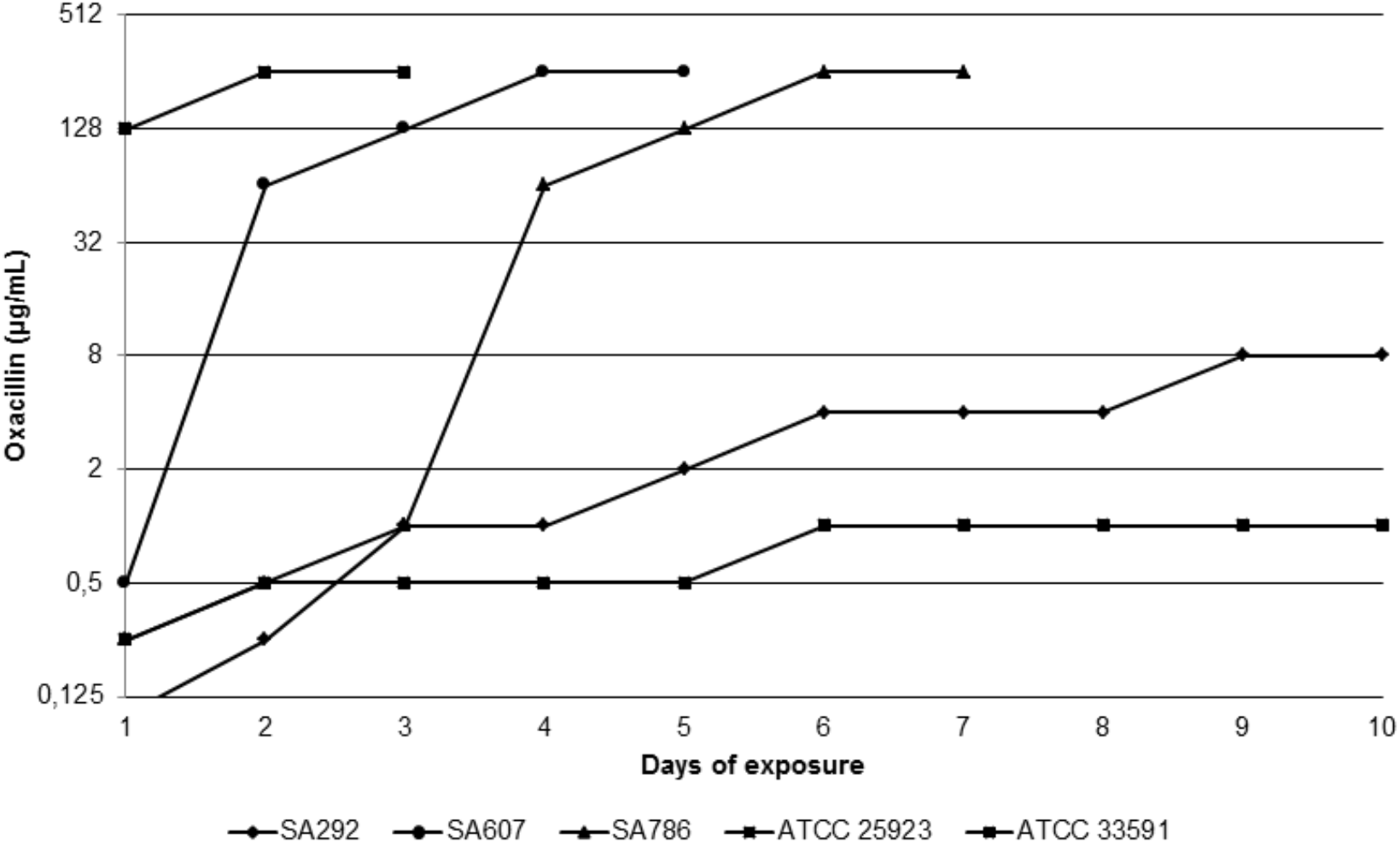
Oxacillin MICs (μg/mL) on days of exposure to oxacillin.

Disc diffusion with cefoxitin confirmed the resistance expression after oxacillin exposure. A reduction of 61% of diameter zone (23.54 mm to 9.17 mm) occurred for SA607. The other OS-MRSA had a reduction of 100% of cefoxitin diameter zone. We also observed changes in measure of oxacillin zone diameter (Fig. 2).

**Figure 2.**
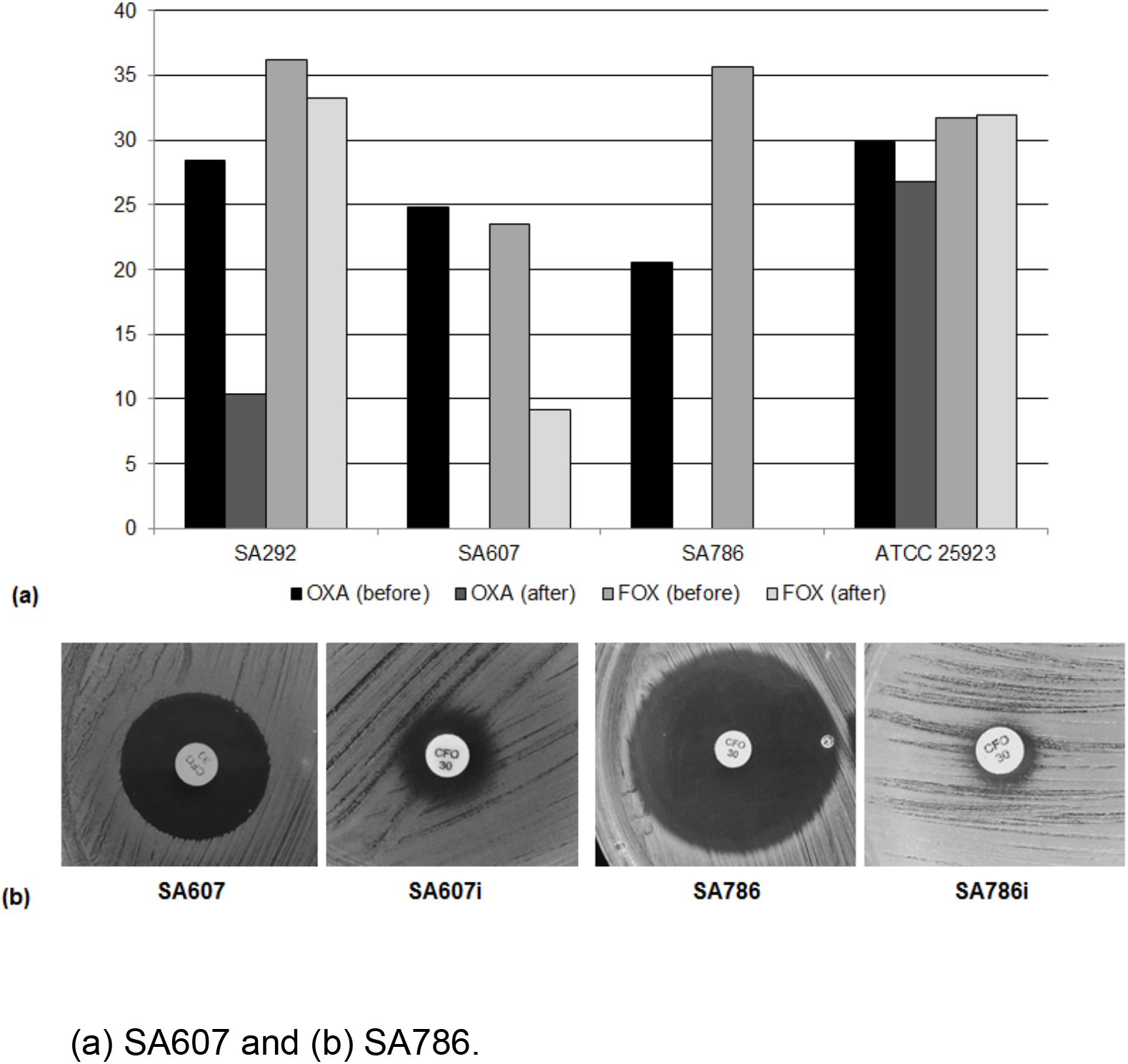
Oxacillin (OXA) and cefoxitin (FOX) zone inhibition diameters (mm) before (left) and after (right) oxacillin exposure.

Some changes in oxacillin MIC and oxacillin zone diameter were also observed for strains used for comparison (SA292 and *S. aureus* ATCC 25923). But in five days the changes were not expressive as noted for OS-MRSA. Therefore, these strains were exposed to oxacillin for 10 days. The β-lactamase positive strain (SA292) had a 64-fold increase, which classify it as oxacillin resistant (MIC > 4 μg/mL). On the other hand, *S. aureus* ATCC 25923 (β-lactamase negative) had 4-fold increase in oxacillin MIC, but continued to display MIC in the susceptibility range (1 μg/mL) (Fig. 1). For both strains cefoxitin zone diameter remained ≥ 22 mm. Thus disc diffusion did not confirm MIC results (Fig. 2).

In addition to changes in susceptibility to oxacillin, both OS-MRSA had notable reduction in penicillin zone diameter, as well as SA292. The reduction of 61.7% of zone diameter (44.42 mm to 17.46 mm) was sufficient to classify SA292i (SA292 induced) as penicillin resistant [15], which make it clear that exposure to oxacillin was able to stimulate β-lactamase production. SA607i and SA786i had 100% reduction in the measurement. After oxacillin exposure no inhibition zone was formed (Fig. 3).

**Figure 3.**
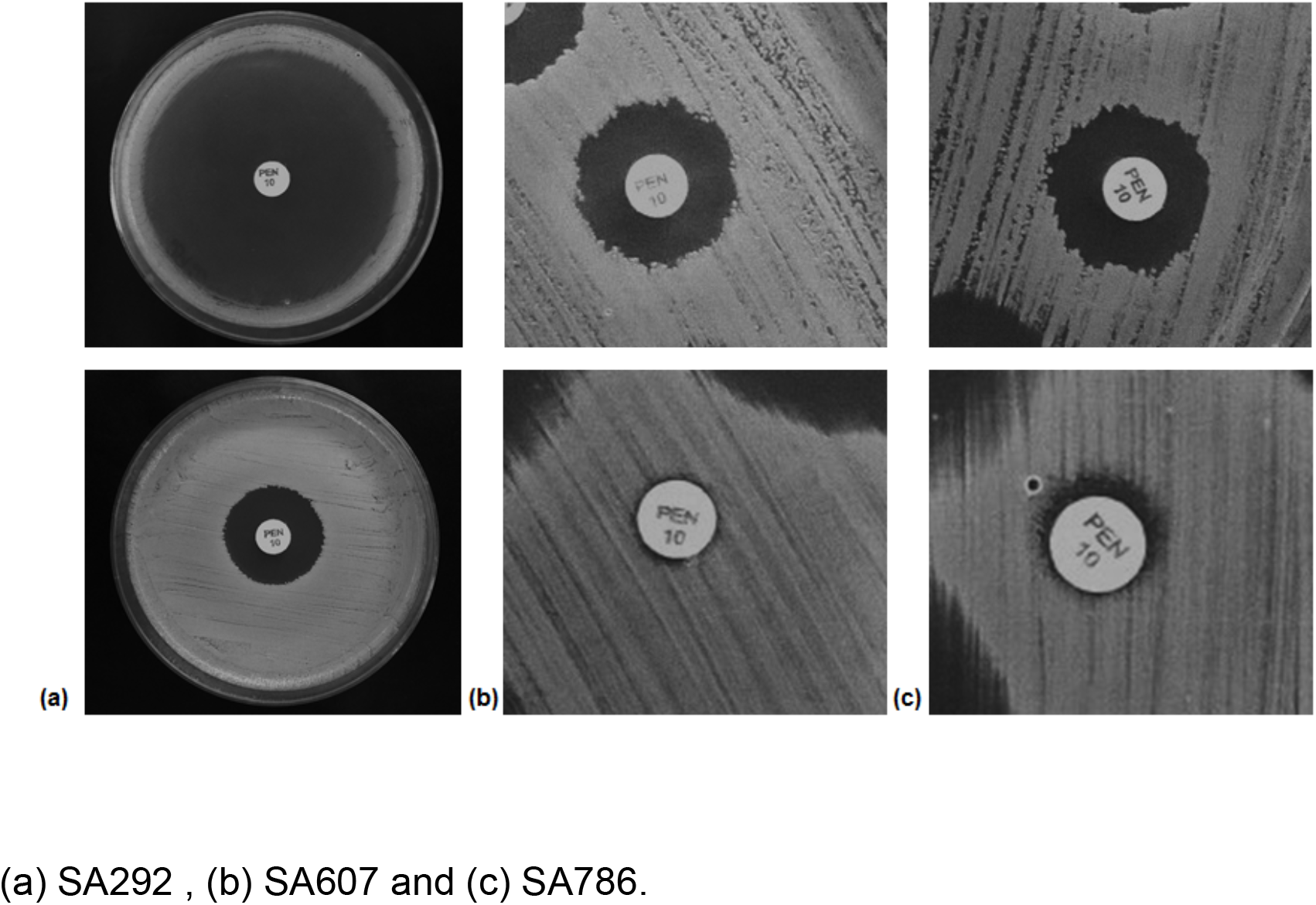
Penicillin zone inhibition of strains before (up) and after (down) exposure to oxacillin.

Different from that was noted for the β-lactam antibiotics (oxacillin, cefoxitin and penicillin), none of the strains altered it susceptibility to non-β-lactam antibiotics.

### Population analysis before and after induction

According to population analysis study, SA607 exhibit oxacillin heteroresistance before exposure to antibiotic (difference between the lowest concentration with maximum inhibition of growth and the highest non-inhibitory concentration > 8-fold). Growth was observed at oxacillin concentrations as high as 32 μg/mL. After oxacillin exposure, growth of 10^4^ to 10^5^ cells was observed at the highest concentration used (256 μg/mL). SA786 was non-heteroresistant, but after oxacillin induction exhibited a result similar to that of SA607 (that is, became homoresistant) (Fig. 4).

**Figure 4.**
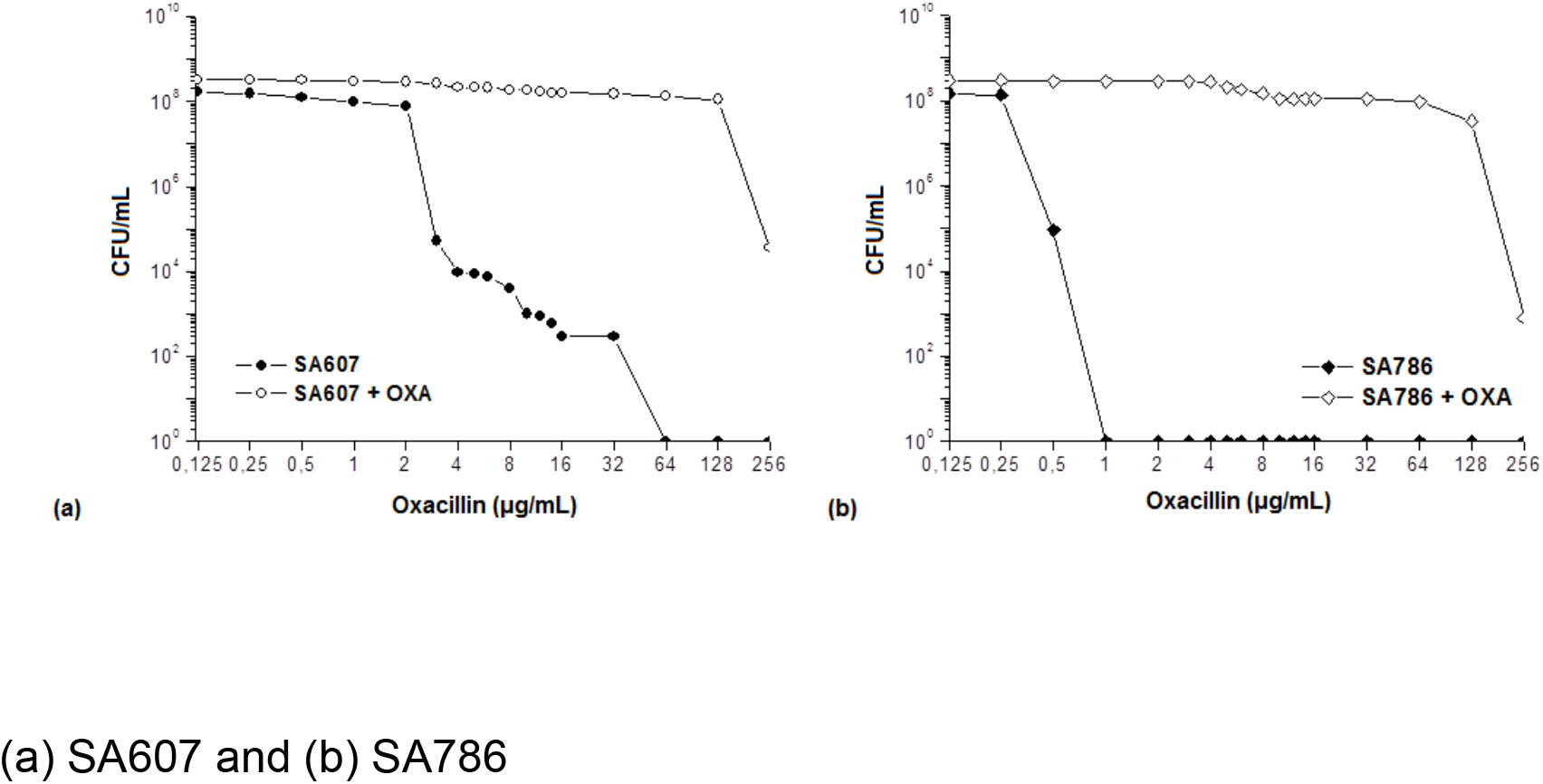
PAP before and after induction with oxacillin. (a) SA607 and (b) SA786

SA292 was defined as homogeneously susceptible (difference ≤ 4-fold), expressing a MSSA curve with no growth at concentrations of oxacillin between 0.5 and 2 μg/mL. This strain remained classified in this way after induction.

## Discussion

In this study, heteroresistant and non-heteroresistant OS-MRSA expressed high phenotypic oxacllin resistance after exposure to sub-MICs of this antibiotic. Thus, both strains were homoresistant (MIC = 256 μg/mL) after *in vitro* antimicrobial selective pressure. Furthermore, we observed that oxacillin sub-MICs resulted in increased β-lactamase production by OS-MRSA.

These results reinforces that the emergence of resistance represents an evolution process in response to antimicrobial selective pressure [22]. Two days were sufficient to cause changes in oxacillin MIC, as previously observed [8–11], but was not able to modify susceptibility to the non-β-lactam antibiotics. Although oxacillin stimulus was maintained for five days and not for long term (more than 10 days), as pointed by Levy & Marshall [23] as a way to select resistant bacteria not only to this antibiotic, but to several others.

We tested only two strains, but our findings corroborate results of other *in vitro* studies [5, 7–12] showing that expression of resistance in OS-MRSA can be altered through stimulation with sub-MICS of β-lactams and suggesting that this kind of *S. aureus* have the potential to express *mecA*-mediated resistance *in vivo*, if exposed to the appropriated antibiotic [6]. Recently, Proulx *et al* [12] and Duarte *et al* [24] related *in vivo* phenotypic resistance conversion in two cases of OS-MRSA bacteremia, one of which was a case of fatal sepsis [24]. Therefore, exposure to β-lactam antibiotics may really result in the emergence of new MRSA strains, as suggested before [4, 13].

Modification of susceptibility profile for oxacillin resistance was observed on the second day of exposure and in four days SA607 was highly resistant (homoresistant by PAP and with oxacillin MIC of 256 μg/mL). Similar results were found by Kampf *et al* [8] for six non-heteroresistant strains. Our non-heteroresistant OS-MRSA (SA786) took longer to change the sensitivity profile – four days to become resistant and seven days to become homoresistant.

Some *in vitro* studies used growth around the cefoxitin disc (in the disc diffusion test) as the inoculum for reexposure OS-MRSA to antibiotic [10, 11]. We highlight that this implies in the selection of subpopulations that survive at higher antibiotic concentrations than a whole heterogeneous population. Therefore, unless OS-MRSA is non-heterogeneous, it is more difficult to determine if changes in expression of resistance are due to the selection of resistant mutants or represent alterations in genic expression or mutations in genes related to resistance.

Although some OS-MRSA are heteroresistant [25] others are truly susceptible [26] (i.e. non-heteroresistant), which does not mean them can not express resistance if exposed to antibiotic [8, 10]. Thus, though heteroresistance represents a natural evolutionary tool for resistance [27], selection of resistant subpopulations is not the only factor to explain the behavior of our OS-MRSA when exposed to antibiotic, because only one of them was heteroresistant. Then we did not rule out the possibility of alterations in *mecA* transcription or mutation/activation of genes that act in bacterial physiology during antibiotic stress [28, 29].

SA786 had no *SCCmec* typable with primers used (types I, II, III and IV), but SA607 had *SCCmec* type IVa, which carries a class B *mec* gene complex. This means that it had truncated (i.e. nonfunctional) *mecR1* gene [30]. Moreover, both strains were resistant to penicillin and therefore have β-lactamase (*bla*) genes. Since the similarity between *mec* and *bla* systems allows that *blaR1* and *blaI* can also regulate the *mecA* transcription [31], we hypothesize that induction of *mecA*-mediated resistance expression after exposure to oxacillin could be related to the action of the β-lactamase (*blaZ*) gene regulators.

This hypothesis was considered because after induction we observed total reduction of penicillin zone diameter in both strains, showing that exposure to oxacillin stimulated β-lactamase production, as a consequence of increased expression of *blaR1* and reduction of *blaI* expression [31]. The change in classification of the susceptibility profile for resistance to penicillin in relation to SA292 made clear that oxacillin induction was able to stimulate β-lactamase production.

*bla* regulators may enhance β-lactam resistance expression in MRSA [32] and we suggest that may have the same effect in OS-MRSA. The expression level of BlaI had already been considered as the mainly responsible for resistance in oxacillin-susceptible *mecA*-positive *Staphylococcus sp.* without *mec* regulators [33]. Furthermore, the involvement of the *bla locus* in the phenotypic expression of oxacillin-resistance in OS-MRSA has already been discussed. Sabat *et al* [26] showed that the inactivation of *blaR1* could be responsible for the susceptibility of a penicillin-susceptible OS-MRSA strain that did not have resistance expression induced when exposed to β-lactams, because when *bla locus* was removed, the strain constitutively expressed the *mecA* gene and therefore oxacillin resistance.

Proulx *et al* [12] believe OS-MRSA strains are mutants in which *mecA* gene is inactive, because its expression in the absence of β-lactam negatively affects its survival. Thus, selection could favor inactivation of the gene in the absence of antibiotics as a way to adapt to the environment. However it is important to take account that the linkage between resistance phenotype and molecular genotype highly varied according to intrinsic resistance, response to antibiotic exposure and presence of genes conferring resistance [34].

*In vitro* induction support this assumption showing that on the other hand, selection may result in adaptation to the antibiotic selective pressure represented by sub-MICs through *mecA* activation, because sub-MICs act on at least three levels: selecting resistant bacteria (pre-existing and new populations), generating genetic and phenotypic variability and acting as signaling molecules, influencing gene expression [2].

Homogeneous resistance expression observed after exposure to oxacillin was related not only to *mecA* activation/expression, but also to β-lactamase production. The results reinforces that OS-MRSA strains have the potential to express β-lactam resistance, even in the absence of resistant subpopulations (i.e. non-heteroresistant strains). This reinforces the impact that irrational use of antibiotics has on individuals colonized by *S. aureus* and on the population, emphasizing that the emergence and spread of resistance to antibiotics represent a process of evolution in response to selective antimicrobial pressure.

However, there are still controversies regarding the influence of selective pressure and the response of OS-MRSA strains to β-lactam antibiotics because little is known about the proportion of strains capable of converting to the resistance phenotype and the genetic mechanisms involved in this conversion [12]. Therefore, further studies are needed to better understand the factors related to induction of resistance expression in the presence of sub-MICs, and whether such induction could also increase the virulence of OS-MRSA, since low concentrations of β-lactam had already been related to increase the expression of virulence factors, such as Panton-Valentine leukocidin, PVL.

## Conclusions

*In vitro* exposure to sub-MICs of oxacillin changed the expression of phenotypic oxacillin resistance in both OS-MRSA studied (one heteroresistant and one non-heteroresistant). Homogeneous resistance expression observed after contact with the antibiotic was related not only to *mecA* activation/ expression, but also to β-lactamse production. The results reinforces that OS-MRSA strains have the potential to express β-lactam resistance, even in the absence of resistant subpopulations (i.e. non-heteroresistant strains).

## Aknowledgments

The authors whish to thank Prof. Natalia Iorio for helpful comments. We also thank to Aline Dias and Fábio Alves for providing OS-MRSA strains.

